# SLAB: A Sweep Line Algorithm in PBWT for Finding Haplotype Block Cores

**DOI:** 10.64898/2026.03.16.712201

**Authors:** Ardalan Naseri, Ahsan Sanaullah, Shaojie Zhang, Degui Zhi

## Abstract

With the increasing availability of high-quality phased haplotype data, researchers can more effectively identify detailed patterns of haplotype sharing and investigate the population genetic processes that shape them. In this work, we define block cores as genomic segments where multiple haplotype blocks overlap. We develop an efficient algorithm to analyze haplotype blocks, focusing on width-maximal matches and the identification of haplotype block cores. We apply our algorithm to UK Biobank haplotypes to quantify block cores and demonstrate the biological and population-level insights that can be inferred from their patterns. Specifically, identified block cores can serve as a basis for detecting signals of selection. Although our results are largely consistent with those from methods such as IBD rate analysis, they also reveal complementary information not captured by IBD rates or other conventional approaches. Source code is available at https://github.com/ZhiGroup/SLAB.

## 1 Introduction

The growing availability of large-scale biobank genomic datasets [1, 2] provides an unprecedented opportunity to deepen our understanding of the relationship between genomic variation and human health, as well as to advance fundamental research in population genetics and evolution. With increasingly high-quality, phased haplotype data (e.g., [3, 4]), researchers can more precisely identify patterns of haplotype sharing and investigate the population-genetic processes that shape them. Identical-by-descent (IBD) segments—long genomic sequences shared between haplotypes that are unlikely to occur by chance—are a powerful indicator of common ancestry. While traditional approaches focus on pairwise IBD between two haplotypes, multi-way IBD sharing is receiving increasing attention [5].

A natural extension of pairwise IBD to multiple haplotypes is the concept of haplotype blocks [6, 7], which represent contiguous genomic regions shared across multiple individuals. The probability that multiple unrelated samples share identical genetic segments by chance is much lower than for a pair of haplotypes, making haplotype blocks particularly useful in population genetics and biomedical research. For example, analyzing shared haplotype blocks can improve the detection of positive selection within populations. These blocks have been shown to effectively identify genomic regions where long, repeated haplotypes suggest recent selective sweeps in large populations [8]. Additionally, haplotype blocks can facilitate the mapping of genetic traits using multi-individual IBD approaches [9]. By combining IBD segments from multiple individuals and adjusting for genome-wide multiple testing, researchers can identify genomic regions associated with complex traits or diseases. Organizing shared genetic variation into coherent blocks enhances statistical power, reduces computational burden, and provides insights into both evolutionary history and genotype–phenotype associations.

We have developed efficient algorithms for identifying haplotype blocks in large cohorts [10, 11]. Given a phased haplotype or genotype panel, these algorithms can efficiently extract all shared segments that exceed a specified length and are present in at least *c* haplotypes. We can efficiently identify blocks that have the maximum number of haplotypes (width-maximal) or the longest genomic/genetic match (length-maximal). For instance, in the UK Biobank, which includes nearly one million haplotypes, finding all width-maximal matches on chromosome 2 (with a length threshold of 2 cM and at least 500 haplotypes) takes less than 2 CPU hours. The algorithms are based on the Positional Burrows-Wheeler Transform (PBWT) [12], which sorts haplotypes by their reversed prefix order. Enumerating all haplotype blocks can thus be achieved in linear time relative to the size of the panel.

However, directly applying these algorithms often results in multiple overlapping blocks that share haplotypes across the same genomic regions. While overlapping blocks may at first appear redundant, the patterns of overlap can be highly informative, potentially revealing underlying population structure, relatedness, or evolutionary processes. For example, overlapping blocks can reflect subgroups of samples that are more closely related and therefore share longer IBD segments in a genomic region. At the same time, smaller blocks within the same region may be preserved across a broader set of haplotypes due to selective pressures, creating a layered structure of overlapping segments. Such patterns capture both recent relatedness and signals of natural selection simultaneously. Yet, such overlapping block patterns are neither well-studied nor well-defined. In this work, we define the block overlapping structure as a *block overlap graph*. We also develop an efficient algorithm to analyze haplotype blocks—focusing on width-maximal matches—and identify the cores of haplotype blocks. We apply our method to UK Biobank haplotypes, quantify block cores, and demonstrate the insights that can be gained from analyzing these block core patterns.

## 2 Methods

### 2.1 Preliminaries

We define a haplotype panel as a binary matrix *X* ∈ {0, 1}^*M*×*N*^, where *M* is the number of haplotype sequences and *N* is the number of variant sites similar to [12]. Each row of *X* represents a haplotype sequence, while each column *X*_*k*_, for *k* = 0, …, *N*− 1, contains the allelic values of all haplotypes at site *k*. Given two haplotypes *p* and *q*, we denote by *p*[*s, e*) and *q*[*s, e*) the subsequences from site *s* to site *e*− 1. A match between *p* and *q* over the interval [*s, e*) is defined when *p*[*s, e*) = *q*[*s, e*).

The concept of haplotype matching can be generalized to *blocks* [11, 13, 14]. *Let C* = {0, …, *M* – 1} be the set of haplotype indices. A *haplotype block* is a tuple *B* = (*H, s, e*), where *H* ⊆*C* is a subset of haplotypes that all share the same subsequence over the interval [*s, e*), with *s < e*. The *length* of the block is *l* = *e* − *s*, and the *width* is *w* = |*H*|. A block is called an (*L, W*)-block if *l* ≥ *L* and *w* ≥ *W* . An *L, W* -block, (*H, s, e*), is called *width-maximal* if the number of matching haplotypes cannot be increased without violating the length constraint. Likewise, an (*L, W*)-block, (*H, s, e*), is *length-maximal* if it cannot be extended without reducing the number of matching haplotypes to fewer than *W* .

In this work, we focus on width-maximal (*L, W*)-blocks and assume the input data is the set of all width-maximal *L, W* -blocks. All width-maximal (*L, W*)-blocks can be detected in *O*(*NM*) using the Positional Burrows-Wheeler Transform (PBWT) [11, 12, 13, 14].

### 2.2 Block overlap graph

The intersection between two haplotype blocks *B*_1_ and *B*_2_ is defined as

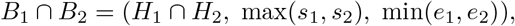

where *H*_*i*_ denotes the set of haplotypes in block *B*_*i*_, and (*s*_*i*_, *e*_*i*_) represent the genomic start and end positions, respectively. The overlap block is considered *non-empty* if *H*_1_ ∩ *H*_2_ ≠∅ and min(*e*_1_, *e*_2_) − max(*s*_1_, *s*_2_) *>* 0.

We further define an (*L, W*)*-overlap* between two blocks if

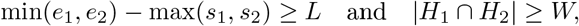

where *L* and *W* are user-specified thresholds for genomic length and haplotype intersection size, respectively. Note that while *B*_1_ and *B*_2_ are maximal blocks, their intersection may not be an (*L, W*)-block.

Based on this definition, we construct a *block overlap graph* (or equivalently, an (*L, W*)-overlap graph), where each node represents a haplotype block and edges connect pairs of blocks that satisfy the (*L, W*)-overlap condition. We identify connected components in this graph, and within each component, we search for the *maximum clique*. The maximum clique, or *core*, represents a maximal set of mutually overlapping blocks that cannot be extended without reducing the overlap degree (i.e., the size of the shared intersection). Thus, the core captures the largest multi-way overlap among a group of haplotype blocks. It is important to note that a maximum clique may not be unique; a connected component can contain multiple maximum cliques of the same size. In such instances, any of these cliques corresponds to a valid block core.

#### Definition: Block core

Let ℬ = {*B*_1_, *B*_2_, …, *B*_*n}*_ be a set of haplotype blocks, where each block *B*_*i*_ = (*H*_*i*_, *s*_*i*_, *e*_*i*_) contains haplotypes *H*_*i*_ and spans the genomic interval [*s*_*i*_, *e*_*i*_].

The block core is defined as the largest subset of blocks that share a common genomic interval and haplotypes. Formally, a subset *C* ⊆ ℬ is *valid* if

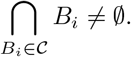

Among all valid subsets, the core block is the one with maximum cardinality:

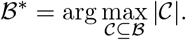

The core interval is the intersection of the intervals of blocks in the core:

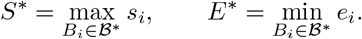

The core haplotype set is the intersection of haplotypes across the blocks in the core:

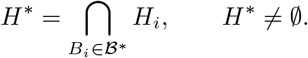

Finally, the block core is represented as the tuple

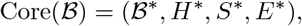

capturing the largest group of blocks that share a common genomic interval and haplotypes.

Figure 1 illustrates a simple example of block cores. In this example, there are two connected components: *B*_1_, *B*_2_, *B*_3_, *B*_4_ and *B*_5_, *B*_6_. The core regions, shown in red, represent the areas where the maximum number of blocks overlap. The figure assumes that the blocks are continuous along the y-axis and can be represented as intervals, although in practice, haplotype blocks may not be strictly continuous along this axis. Notably, blocks *B*_4_ and *B*_1_ also overlap, but with only two overlapping blocks; hence, this region forms a *maximal clique* rather than a core.

**Figure 1:**
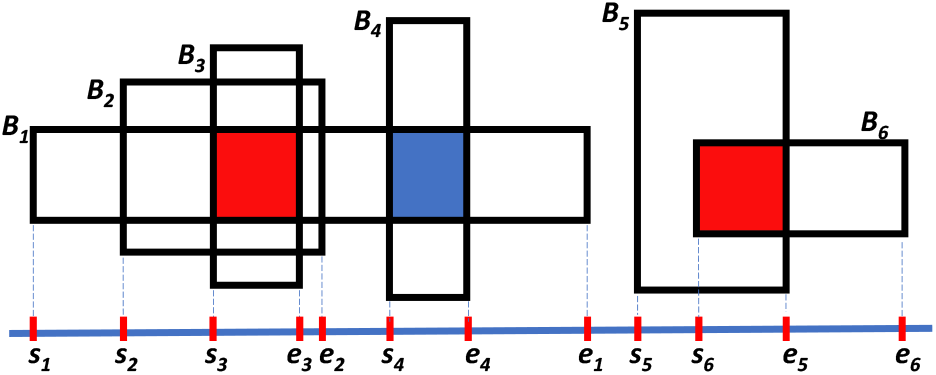
Schematic illustration of block cores. The example contains two connected components, with core regions (maximum cliques) highlighted in red. The overlap between the blocks *B*_1_ and *B*_4_, highlighted in blue, is a maximal clique.

### 2.3 Local IBD graph (LIG) and local IBD graph sequence (LIGS)

To relate block overlaps to classical representations of shared ancestry, we interpret them using the framework of local IBD (identity-by-descent) graphs. An IBD segment between two haplotypes, *h*_*i*_ and *h*_*j*_, is defined as a continuous genomic interval [*s, e*) where both haplotypes are identical by descent. This means they share a common ancestral haplotype without any recombination occurring within the interval [*s, e*).

#### Local IBD graph (LIG)

At any genomic position *k*, we define the *Local IBD graph (LIG)* as

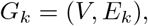

where *V* is the set of haplotypes (or samples) and *E*_*k*_ is the set of edges connecting haplotypes that share IBD at site *k*. In other words, an edge (*i, j*) ∈ *E*_*k*_ exists if haplotypes *h*_*i*_ and *h*_*j*_ share an IBD segment covering site *k*.

If we restrict this definition to IBD segments of at least length *L*, we define the *L-IBD Graph* at site *k*, denoted 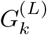, as the subgraph including only those IBD edges whose segments span at least *L* units:

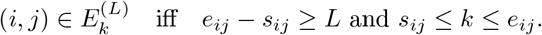

where *L* may refer to a physical or a genetic length threshold or the number of variant sites.

#### W-cliques

A set of at least *W* haplotypes 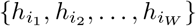 that all share IBD at site *k* forms a *W -clique* in the Local IBD Graph (LIG) *G*_*k*_. For all distinct pairs 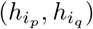 within the set, (*i*_*p*_, *i*_*q*_) ∈ *E*_*k*_. We refer to such structures as *W -cliques in the LIG*. These cliques represent multi-way IBD sharing events among haplotypes at a specific genomic site and serve as indicators of localized shared ancestry within a population.

#### Local IBD graph sequence (LIGS)

The local IBD graph at site *k* is typically highly similar to that at the adjacent site *k* + 1. Capturing haplotype sharing that persists across contiguous sites provides a more coherent view of local genetic relationships along the genome. In this context, *W* -cliques— cliques that remain intact across a range of consecutive sites among multiple individuals—represent stable substructures within the local IBD graph sequence. Tracking these persistent cliques enables the identification of genomic intervals where shared haplotypic relationships remain consistent. We refer to the set of sites over which a clique persists as its *persistence range*. Note that a *L*-persistent *W*− clique in LIGS is a (*L, W*)-block. The reason we introduce the LIG is to give exact locality to overlapping blocks, which is useful for gene mapping.

#### Local cores in block overlap graph

We define local cores as maximal cliques in the block-overlap graph. A local core is a set of blocks that all overlap with one another and cannot be extended—adding any additional block would break this all-overlapping property. A connected component may contain multiple local cores, each representing a maximal set under this definition. In the example shown in Figure 2, the graph contains two connected components, within which six local cores, *c*_1_, …, *c*_6_, are identified. Among these, the two largest cores, *c*_5_ and *c*_6_, are maximum cliques (cores) and are also included as part of local cores.

**Figure 2:**
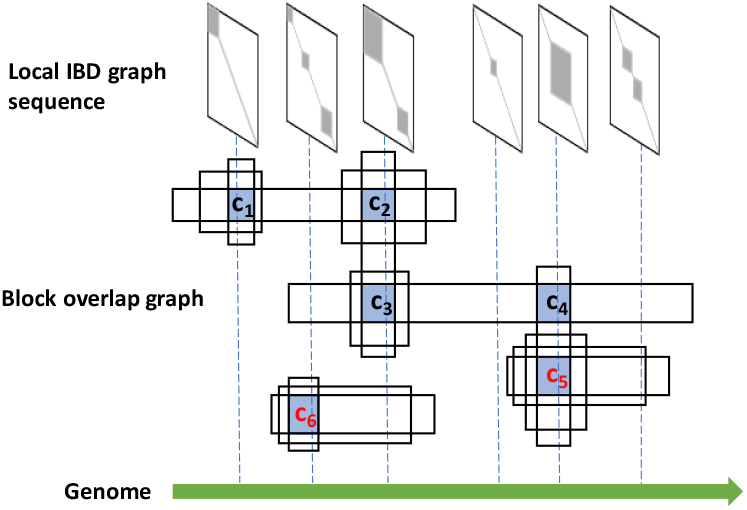
Schematic of the block-overlap graph and the corresponding local IBD graph sequence. Local cores, defined as maximal sets of blocks that all overlap, are identified from the block-overlap graph and are highlighted in blue. *c*_5_ and *c*_6_ are two haplotype block cores, the intersections of the maximum number of blocks in two different connected components.

#### Local core graph

Local cores and their relationships can be represented using a core graph, where each core is represented as a node, and edges connect cores that share one or more blocks. This representation is primarily intended for exploratory analysis of relationships among detected cores and is not required for the core detection algorithm itself. Large cores with blocks that have similar boundaries may point to redundant information, which can be quantified using intersection-over-union (IOU) measures over haplotypes, IOU(*W*), or genomic ranges, IOU(*L*). For cores with many blocks, median-based measures such as median(*W*)*/* max(*W*) and median(*L*)*/* max(*L*) may provide a more robust assessment.

#### Terminology

In the block overlap graph, a maximum clique corresponds to a block core, while maximal cliques correspond to local cores. A maximum clique (core) is the largest clique in a component, whereas a maximal clique (local core) cannot be extended further but is not necessarily the largest.

### 2.4 SLAB framework for block overlap graph analysis

We developed a set of algorithms to characterize structural patterns within the block overlap graph. These algorithms aim to uncover highly connected substructures—such as block cores and persistent cliques. Given user-defined thresholds for block definition (*L*_block_, *W*_block_), we first identify all width-maximal haplotype blocks. Using additional thresholds for defining block overlaps (*L*_overlap_, *W*_overlap_), we construct the block overlap graph, where nodes represent haplotype blocks and edges indicate pairwise overlaps that satisfy these criteria. Within each connected component of this graph, we identify the *maximum clique*—the largest set of mutually overlapping blocks—which we refer to as the *block core*.

#### 2.4.1 A Sweep Line algorithm for finding block cores (maximum cliques)

Given a set of blocks *B*_1_, *B*_2_, …, *B*_*n*_, where each block is represented as (*H*_*i*_, *s*_*i*_, *e*_*i*_), our goal is to find the largest subset of blocks in which all blocks overlap. This problem is analogous to the *maximum clique problem*, which is known to be NP-hard. Our formulation can be reduced to the classical maximum clique problem by representing each block as a node and connecting two nodes when the corresponding blocks overlap. Finding the largest overlapping subset of blocks is therefore equivalent to finding the maximum clique in this graph. However, unlike the general clique problem, haplotype blocks are ordered by their genomic start and end positions. This ordering, together with the positional structure of haplotypes in the positional Burrows–Wheeler transform (PBWT), can be exploited to improve computational efficiency and allow an efficient polynomial-time solution under the structural constraints induced by genomic ordering and PBWT rank properties.

To achieve polynomial time complexity in finding block cores, sorting plays a key role in enabling efficient block comparisons. We adapt a two-dimensional version of the sweep line algorithm to identify the maximum clique. Each block generates two events (a start and an end), which are sorted and processed sequentially. When an end event is reached, it may mark the point at which the maximum clique occurs in the block ending at that position. At this stage, we compare the haplotypes within the overlapping blocks.

This step relies on the rank of haplotypes, defined as the inverse of the positional prefix array (PPA) in the PBWT. The PPA represents the order of haplotypes in the PBWT matrix, while its inverse gives the rank of each haplotype. We store the inverse PPA for all sites in the PBWT. During the sweep, we maintain a set of active blocks, and for each active block, we compute the rank of its haplotypes at the current end event. The relative order of haplotypes in the positional prefix array remains invariant from the starting to the ending site of a block. Therefore, a divide-and-conquer algorithm can be applied to find the range (interval) for each haplotype block. The sweep line algorithm then identifies regions of maximum overlap, enabling us to efficiently detect the maximum clique. Figure 3 illustrates a schematic of the approach. Algorithms 1 and 2 in Appendix A describe the overall procedure and the subroutine used to compute allowable intervals based on haplotype ranks, respectively. 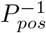 gives the rank of haplotypes in reverse prefix order at the site *pos*, and *AdjList* encodes the adjacency structure of blocks (i.e., which pairs of blocks overlap). *Component* contains the block IDs belonging to a connected component, and *Blocks* stores all blocks, each with a starting site, an ending site, and the associated haplotype list *H*.

**Figure 3:**
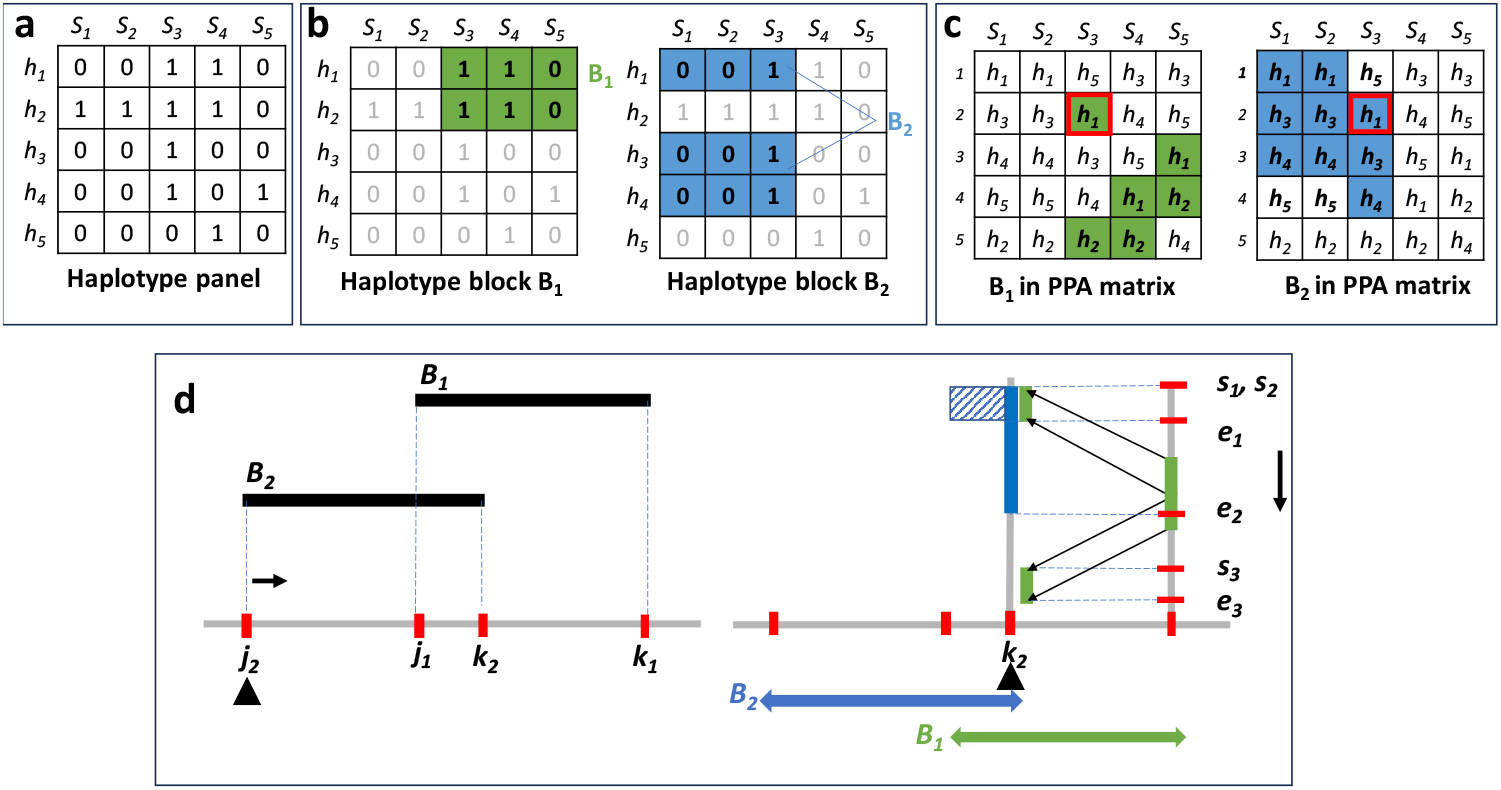
Schematic illustration of the algorithm for finding maximum overlapping blocks in PBWT. This example contains a haplotype panel containing 5 haplotypes and 5 variant sites (a). Haplotype blocks *B*1 and *B*2 are highlighted (b). Haplotypes at the end of each block are adjacent to each other, but along the match may be interrupted by other haplotypes. *h*3 and *h*4 are between the matching block containing *h*1 and *h*2 (c). Finding maximum overlapping blocks (cores) is achieved using a 2-D sweep line algorithm. Each event corresponds to the start or end of a block. At the block’s ending site *k*_2_, the haplotypes in other block(s) in the active set are projected onto the current site, and the sweep line algorithm is applied along the y-axis to identify the maximum overlapping blocks (intervals) using the projected intervals.

#### Time complexity

Let *B* denote the total number of haplotype blocks. During the sweep-line procedure, blocks whose genomic intervals overlap the current sweep position are maintained in an active set; let *a* denote the maximum number of active blocks at an event. Each active block is projected onto the positional prefix array (PPA), producing contiguous rank ranges that define disjoint intervals along the y-axis. Let *i* denote the maximum number of such rank intervals examined during overlap comparison. The algorithm processes 2*B* start and end events, which are sorted in *O*(*B* log *B*) time. Since interval comparisons among active blocks are performed at each ending event, the overall time complexity is

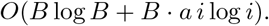

#### 2.4.2 Persistent maximum cliques

To enforce a minimum core length *L*, we modify the sweep line procedure as follows. For each *ending event*, corresponding to the ending site *e* of a block under consideration, we define the subset of active blocks to include only those whose span satisfies the length threshold:

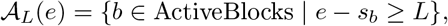

where *s*_*b*_ is the starting site of block *b*. In other words, only blocks whose length from start to the current ending site is at least *L* are considered when forming cliques. This ensures that all resulting cores have a minimum length of *L*, measured in genomic coordinates, genetic distance, or number of sites. A clique spanning over *L* units is analogous to *persistent ranges* in LIGS.

#### 2.4.3 A Sweep line algorithm for finding local Cores (maximal cliques)

Similar to the algorithm for finding the maximum core, we employ a 2-D sweep line algorithm to identify all maximal cliques. In this approach, the triggering events correspond to the start and end of each block, with each block generating two events. The events are first sorted, and as we iterate through them, a start event adds the block to the active list, while an end event projects all haplotypes from the active blocks to the current position of the event. The resulting intervals along the y-axis define candidate cliques: for each interval, the set of overlapping blocks in the active list is enumerated and considered as a candidate clique.

Unlike the maximum core algorithm, here we must track all candidate cliques and validate each one to ensure maximality. At each ending event, all combinations of overlapping blocks observed in the intervals are collected. The candidate sets are then sorted by size, and processing begins with the largest sets, retaining only maximal cliques; for example, if *B*_1_, *B*_2_, *B*_3_ form a maximal clique, any subsets such as *B*_1_, *B*_2_ or *B*_1_ that appear among the candidates are discarded. Algorithm 3 in Appendix A provides a description of the procedure for finding local cores.

## 3 Results

### 3.1 Run time and memory usage

We initially employed the P-smoother [10] to correct genotyping errors, followed by the c-PBWT (forward PBWT [15]) algorithm to identify haplotype blocks containing at least 500 haplotypes over distances of 1 and 2 cM, using phased data from the UK Biobank (UKBB) [1]. The PPA matrix was loaded into memory, resulting in a maximum memory usage of 278 GB for the longest chromosome (chr1). This high memory usage is primarily due to the large size of the matrix, which is nearly 194 GB. To reduce memory consumption, a more efficient implementation could be considered, such as using a memory-mapped version or loading portions of the array at a time.

Our implementation first detects connected components within the haplotype blocks based on user-defined parameters, including the number of samples and the length of shared blocks. We then identify the cores of each connected component that exhibit the maximum number of overlaps. We computed connected components among haplotype blocks in the UK Biobank. Two blocks are considered connected if they share at least 500 haplotypes over distances of 1 or 2 cM, respectively.

Afterward, our method searches for the core within each connected component. The total processing time for all autosomes, which includes identifying connected components and maximum cliques, was about 4 hours for 2 cM blocks and 32 hours for 1 cM blocks, utilizing an AMD EPYC 7763 64-Core processor. The computational runtime across all autosomes of the UK Biobank dataset is shown in Figure 4. There are significantly more blocks with the 1 cM cut-off, which explains why the runtime for 1 cM is nearly eight times longer.

**Figure 4:**
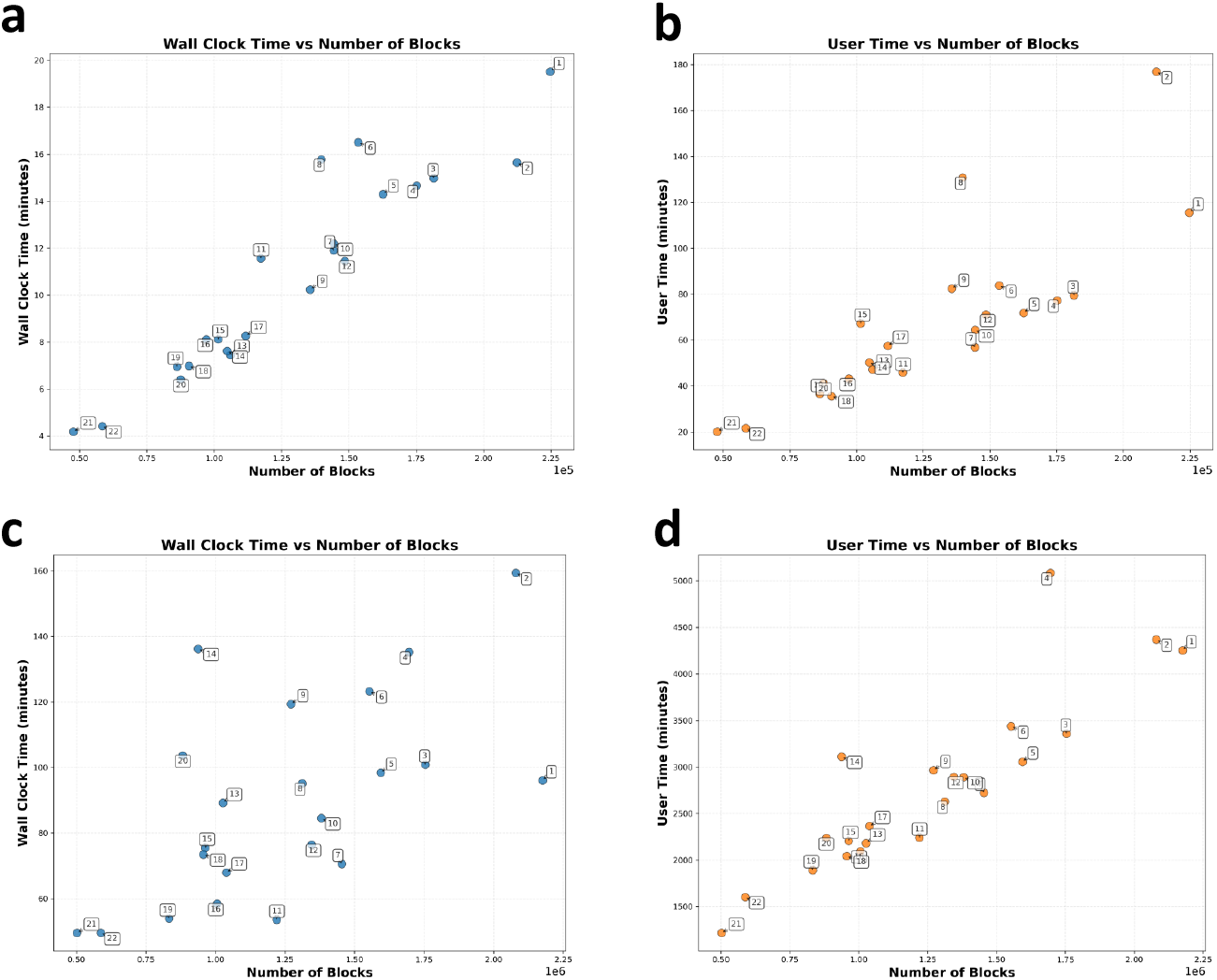
Run times for identifying connected components and cores across all autosomes in the UK Biobank. Panels (a) and (b) show results for 2 cM blocks, while (c) and (d) for 1 cM blocks.

### 3.2 Haplotype block cores in real data (UKBB)

In this section, we analyze genome-wide maximum cliques (block cores) and locally examine maximal cliques (local cores) in selected regions. A total of 168,872 cliques (cores) were found out of 2,831,125 blocks (*L*=2 cM). The largest core contained 980 blocks, while the average maximum clique size was 13. The largest clique identified across the autosomes in the UK Biobank is located on chromosome 6. This clique includes blocks that range from positions 23,927,186 to 32,681,645 (chr6/hg19). This region covers the extended major histocompatibility complex (xMHC) [16], with 980 blocks located at chr6:25,414,023–32,681,645. Notably, the number of haplotypes shared across all blocks in the maximum clique (Intersection Samples) is not the largest observed along the chromosome (see Figure 5).

**Figure 5:**
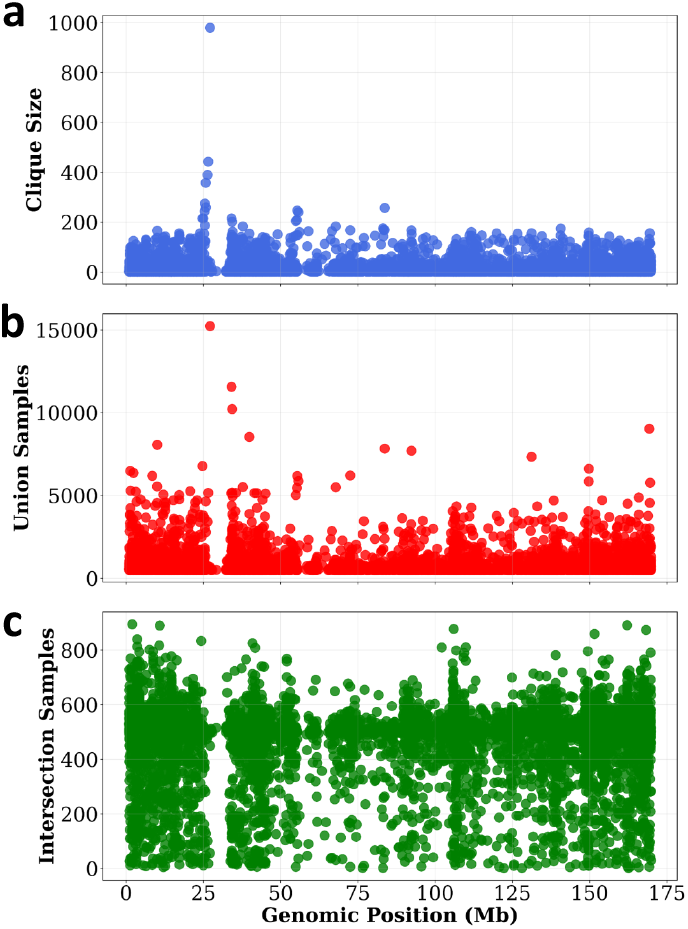
The extended MHC region in chromosome 6 contains the largest maximum clique. The x-axis shows the genomic position corresponding to the midpoint of the haplotype block cores.

The second-largest clique is located on chromosome 3, spanning genomic regions from 43,201,732 to 53,535,073 (p21.31, hg19). Within this region, 341 blocks overlap the gene *SLC6A20*. A Neanderthal-derived haplotype within this gene has been reported as a major risk factor for severe COVID-19 [17]. According to Zeberg and Pääbo [17], the genetic variants most strongly associated with severe COVID-19 on chromosome 3 (45,859,651–45,909,024, hg19) are in high linkage disequilibrium (LD). These haplotypes are believed to have been introgressed into modern humans from Neanderthals or Denisovans approximately 40,000–60,000 years ago [18, 17].

We analyzed all individuals within the detected core, along with the rest of the cohort, using their whole-genome sequencing data [19]. Specifically, we examined the variant sites reported in [17] as part of the haplotype risk factor. For this analysis, we used Data-Field 24308 (DRAGEN population-level WGS variants, PLINK format, [500k release]) available in UKB-RAP, and computed the allele frequencies for individuals within the block core and for the remainder of the cohort. As shown in Table 1, individuals within the haplotype blocks of the maximum clique exhibit approximately half the allele frequency observed in the rest of the cohort, suggesting a lack of the haplotype carrying the risk alleles.

**Table 1:** Allele frequencies among samples within haplotype blocks corresponding to the maximum clique on chromosome 3, compared with the remaining cohort (hg38 coordinates). The coordinates have been lifted over from hg19 to hg38 from the reported loci in [17].

| Chr | Position | SNP ID | REF | ALT/Risk | AF (max clique) | AF (rest of cohort) |
| --- | --- | --- | --- | --- | --- | --- |
| 3 | 45867532 | rs35044562 | A | G | 0.0353 | 0.0736 |
| 3 | 45859597 | rs73064425 | C | T | 0.0353 | 0.0734 |
| 3 | 45858159 | rs34326463 | A | G | 0.0353 | 0.0736 |
| 3 | 45866624 | rs13081482 | A | T | 0.0351 | 0.0730 |
| 3 | 45838989 | rs35508621 | T | C | 0.0353 | 0.0736 |
| 3 | 45823240 | rs10490770 | T | C | 0.0353 | 0.0736 |
| 3 | 45821460 | rs71325088 | T | C | 0.0352 | 0.0735 |
| 3 | 45820440 | rs13078854 | G | A | 0.0352 | 0.0734 |
| 3 | 45818159 | rs17713054 | G | A | 0.0352 | 0.0735 |
| 3 | 45830416 | rs67959919 | G | A | 0.0352 | 0.0735 |
| 3 | 45847198 | rs34288077 | A | G | 0.0353 | 0.0736 |
| 3 | 45848429 | rs35081325 | A | T | 0.0353 | 0.0736 |
| 3 | 45825948 | rs35624553 | A | G | 0.0353 | 0.0736 |

We also computed IBD rates using hap-IBD [20] (see Figure 6g and h) with a window size of 100k and a target IBD length of 2 cM. On chromosome 2, the genomic span of blocks in the maximum clique coincides with a peak in the IBD rate. The highest IBD rate region (0.31%) on chromosome 2 includes the gene *LCT*, which carries a variant associated with lactose tolerance in Europeans [21]. In contrast, the IBD rate peaks on chromosome 3 do not overlap with the identified maximum clique.

**Figure 6:**
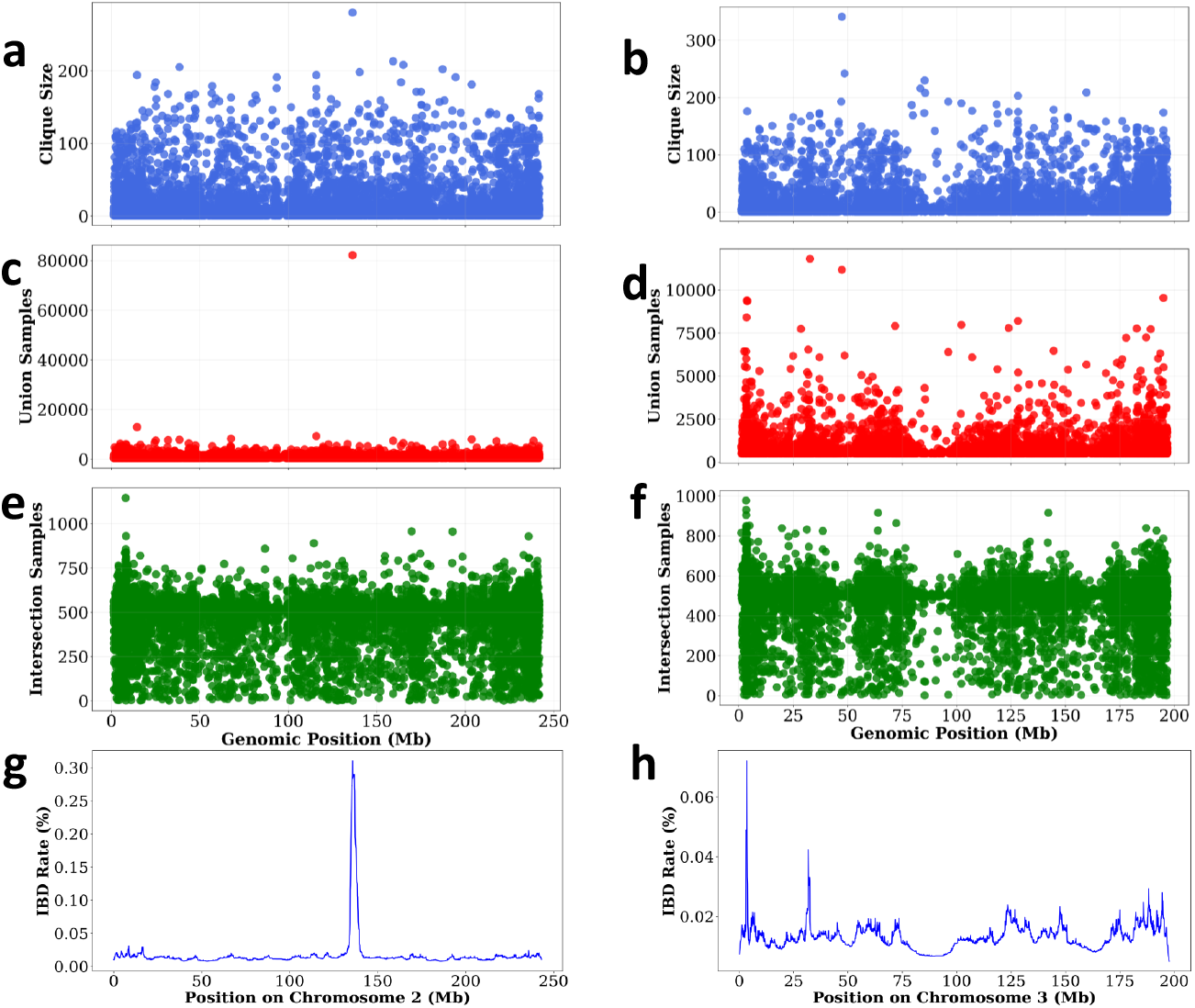
Maximum cliques highlight potential signals of natural selection. On chromosome 2, there is a strong signal where a large number of haplotypes share the same sequence. In contrast, chromosome 3 shows a prominent peak in clique size despite involving fewer shared haplotypes, suggesting a possible selection signal not captured solely by IBD rate.

We also applied the maximal clique method to UK Biobank chromosome 22 using haplotype blocks of length 2 cM (58,426 blocks) and a minimum local core length of 1.5 cM. Each core contained, on average, approximately 17 blocks. The analysis ran for 2 hours and 22 minutes and identified 13,325 maximal cliques. On chromosome 2, we selected all blocks overlapping genomic position 135,755,899, corresponding to the location of the largest clique. In this region, there were 7,767 blocks, and we detected 3,658 maximal cliques (local cores). The number of cliques in this region is notably higher than that observed for chromosome 22, likely reflecting the increased complexity of the local haplotype structure.

Table 2 illustrates the ethnic-background composition of the union and intersection of haplotypes for an example clique of size 37. This clique was selected because it was among the limited number of cliques that contained more than 10% of individuals with ethnic backgrounds other than British or Irish. In the union, most individuals are British, with a small percentage being Ashkenazi Jewish. In the intersection, the British count drops, and at least 19 of the remaining haplotypes are Ashkenazi Jewish. Ethnic background information was obtained from self-reported metadata in UK Biobank (Data-Field 21000), and Ashkenazi Jewish ancestry was assigned using the method described in [22]. Note that the union and intersection counts are for haplotypes, and a single haplotype can have multiple ethnic backgrounds assigned.

**Table 2:** Ethnic-background counts for the union and intersection of haplotypes in a maximal clique on chromosome 2 (clique size = 37; union = 1,147; intersection = 258).

| Ethnic background | Union count | Intersection count |
| --- | --- | --- |
| British | 1081 | 241 |
| Any other White background | 31 | 13 |
| Irish | 22 | 3 |
| Ashkenazi | 22 | 19 |
| Any other mixed background | 2 | 0 |
| Caribbean | 2 | 0 |
| Other ethnic group | 2 | 1 |
| White and Black Caribbean | 1 | 0 |
| White and Asian | 1 | 0 |
| White | 1 | 0 |

## 4 Discussion and Conclusions

We present a novel and efficient method to compare haplotype blocks and identify their core regions. Our approach leverages the positional ordering of haplotypes and a two-dimensional sweep line algorithm to detect overlapping structures among blocks. By combining sorting and rank-based comparisons, we efficiently determine shared core regions across groups of overlapping haplotype blocks. This enables scalable analysis even in large genomic datasets, where the number of potential overlaps can be substantial. The computational complexity of the method is *O*(*B* log *B* + *B* · *a i* log *i*), where *B* is the total number of blocks, *a* is the number of active blocks, and *i* is the number of intervals used when comparing haplotypes. In practice, the number of intervals along the y-axis (*i*) is typically much smaller than the number of haplotypes within each block, although the worst-case scenario scales with the sample size. Despite this, our method remains highly efficient: processing a 2-cM genomic region typically requires only a few minutes per chromosome in the UK Biobank, which contains almost 1 million haplotypes. The proposed framework can be further extended to merge haplotype blocks efficiently. By tracking overlapping and adjacent intervals, it is possible to consolidate blocks that share substantial haplotype content or overlap beyond a specified threshold.

We note that IBD blocks and cores are closely related to the concept of IBD graphs, which provide a powerful framework for defining communities in biobanks [23, 24, 25]. Traditional IBD graphs are defined globally, with each individual as a node and edges representing shared IBD segments. However, such global graphs do not capture the genomic locality of IBD sharing. In the Methods, we introduced local IBD graphs and local IBD graph sequences, which better reflect the structure of shared segments along the genome. A local IBD graph for a genomic region connects individuals who share IBD segments within that region. Features derived from these graphs, such as the IBD rate—the fraction of individual pairs sharing a segment—can reveal demographic history and selection.

Local IBD graphs also facilitate the identification of sub-populations sharing specific segments, highlighting independent selection signals in regions such as *LCT*. Nearby local IBD graphs are often highly similar, leading to redundancy in representation. Treating them as a sequence, defined as local IBD graph sequences in the Methods, allows efficient representation and analysis, inspired by tree sequences from ARG studies. Nevertheless, the explicit construction of the entire local IBD graph sequence can be daunting in large biobanks. Our blocks capture large cliques that persist across multiple local IBD graphs at adjacent sites, while local cores represent persistent, dense clusters. Both blocks and local cores provide less-redundant representations of local IBD graph sequences, enabling more efficient analysis and deeper insights.

The term “core haplotype” has previously been used to refer to a locus of interest. The age of such a core haplotype has been studied using extended haplotype homozygosity (EHH), a measure of signatures of positive selection [26]. EHH is defined as the probability that two randomly chosen chromosomes carrying the core haplotype remain identical from the core region to a given point along the chromosome. Intuitively, EHH measures how far haplotypes remain identical as one moves away from a core site. Under positive selection, high levels of haplotype homozygosity extend much farther than expected under neutrality; to capture this effect, Voight et al. [27] proposed integrating the observed decay of EHH away from a specified core allele. This quantity, known as integrated haplotype homozygosity (iHH), is computed until EHH falls below a predefined threshold. Both EHH and iHH were formulated in terms of homozygosity, reflecting their development during an era when genotype data were predominantly unphased. Although our term “block core” may appear similar to the haplotype core of [26, 27], the two concepts differ substantially. Those earlier definitions treat the core as an arbitrary locus or region characterized by high allele frequency. In our framework, by contrast, the core is not chosen arbitrarily; instead, it emerges from minimum thresholds on haplotype length and the number of samples supporting it, which together determine the resulting structure. A further distinction is that we work with phased data and track haplotype identity directly, rather than relying on homozygosity as a proxy. Finally, our block core definition is designed to scale to biobank-sized datasets, whereas the original haplotype core methods were developed for and applied to cohorts far smaller than those available today.

In this work, we did not jointly analyze the clique size and the intersection size of samples in overlapping blocks. At genomic positions where the maximum clique is observed, the intersection of samples across overlapping blocks may not be maximal. A small intersection combined with a large clique may indicate haplotype variation surrounding a shared core. Within the Extended Haplotype Homozygosity (EHH) framework, stronger signals are associated with longer-range homozygosity and would typically produce larger sample intersections. Therefore, a large maximum clique size accompanied by a relatively small intersection may reflect population-specific selection or differences in genetic background. Although the union of samples in overlapping blocks is expected to be largest near the position of the maximum clique, it does not necessarily coincide with it. In our real data analyses, we focused primarily on identifying and characterizing maximum cliques, highlighting representative examples. Further work could uncover more complex evolutionary patterns, such as recombination hotspots or population-specific haplotype structures, using local cores. Beyond population genetics, this framework has potential applications in genome-wide association studies (GWAS), where identifying shared haplotype cores may help localize causal variants and refine association signals.

The blocks used as input in our results were defined with a minimum length in cM. A constraint on the minimum number of sites can be easily incorporated into the block detection step, thereby avoiding potential artifacts arising from low-density marker regions. In our results, the detected peaks were not necessarily located near low-density regions (e.g., centromeres), but this consideration may be important for other chromosomes or datasets. We also did not investigate the effect of phasing errors. Phasing quality improves substantially once sample sizes exceed 50,000 individuals [4]. Data sets such as the UK Biobank therefore achieve very high phasing accuracy due to their large sample sizes. We do not expect that phasing errors substantially affect results in large cohorts. In panels containing hundreds of thousands to millions of haplotypes, phasing errors, especially for shorter segments (e.g., 0.5 cM), are expected to be negligible. Nevertheless, further investigation in future work would be valuable.

## 5 Acknowledgements

This research has been conducted using the UK Biobank Resource under Application Number 24247. This work was supported by the National Institute on Aging of the National Institutes of Health under Award Number R01AG081398 and the National Human Genome Research Institute under Award Number R01HG010086.

## Appendix A

### Algorithm 1

Find maximum clique (block core)

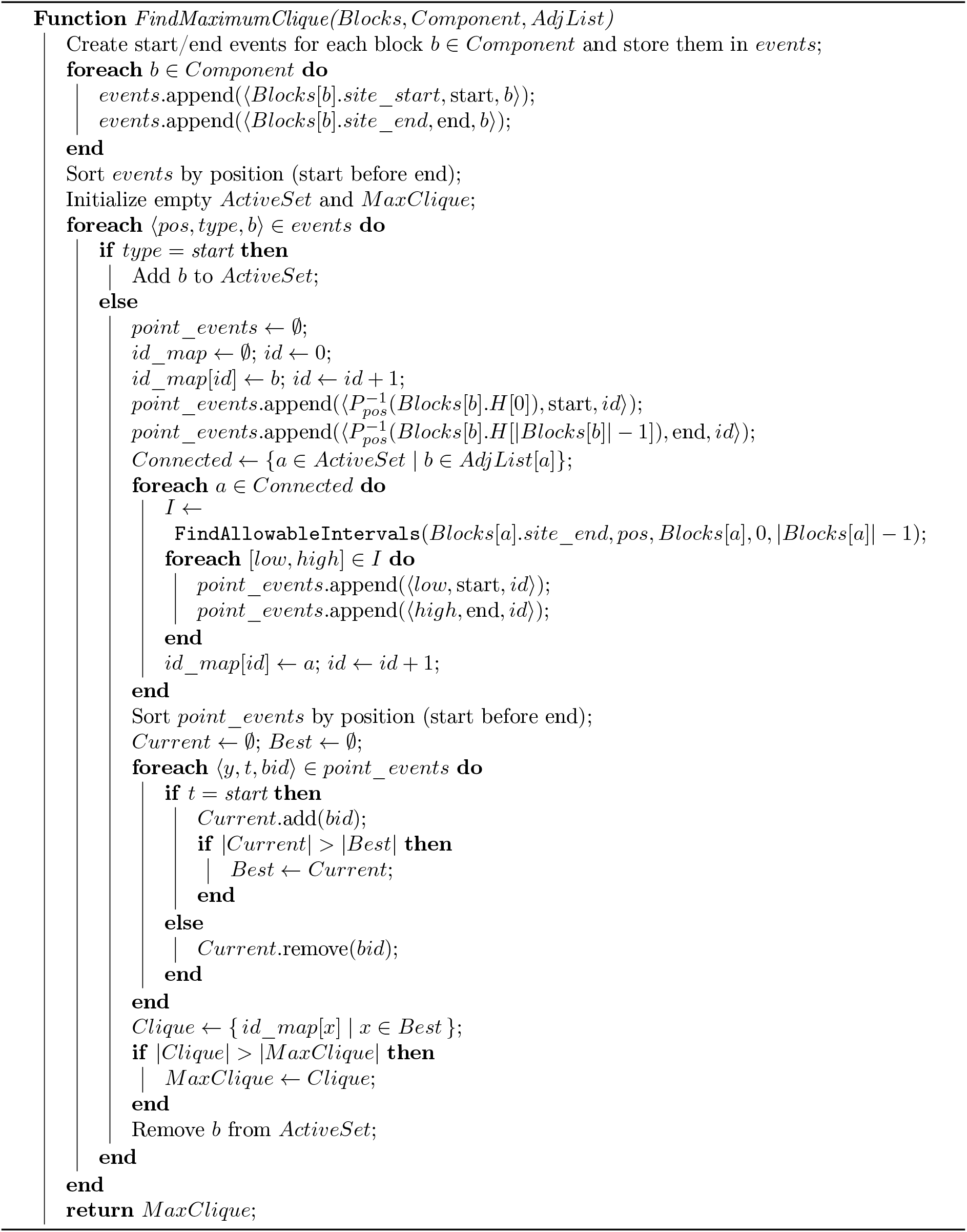

### Algorithm 2

Find allowable intervals.

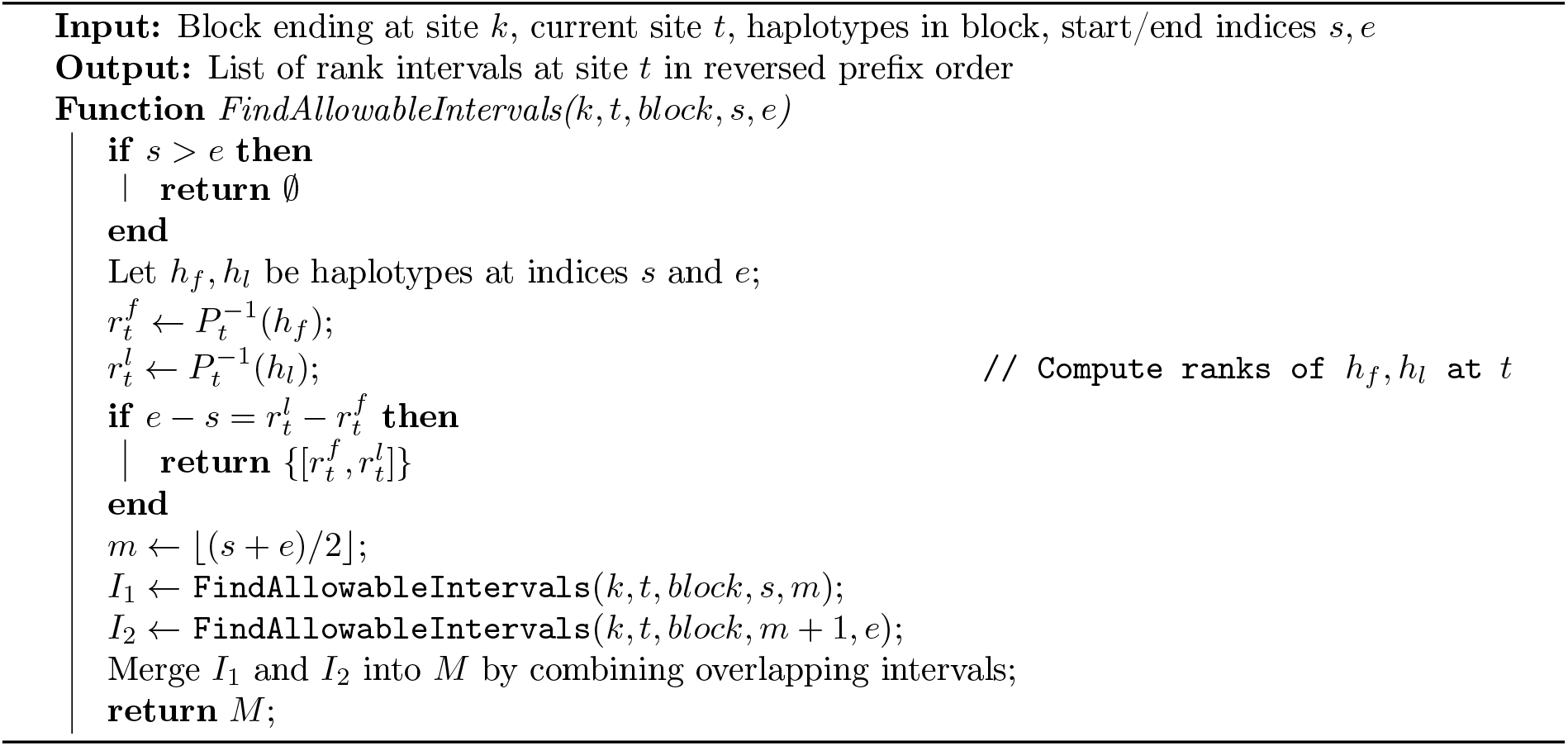

### Algorithm 3

Find local cores

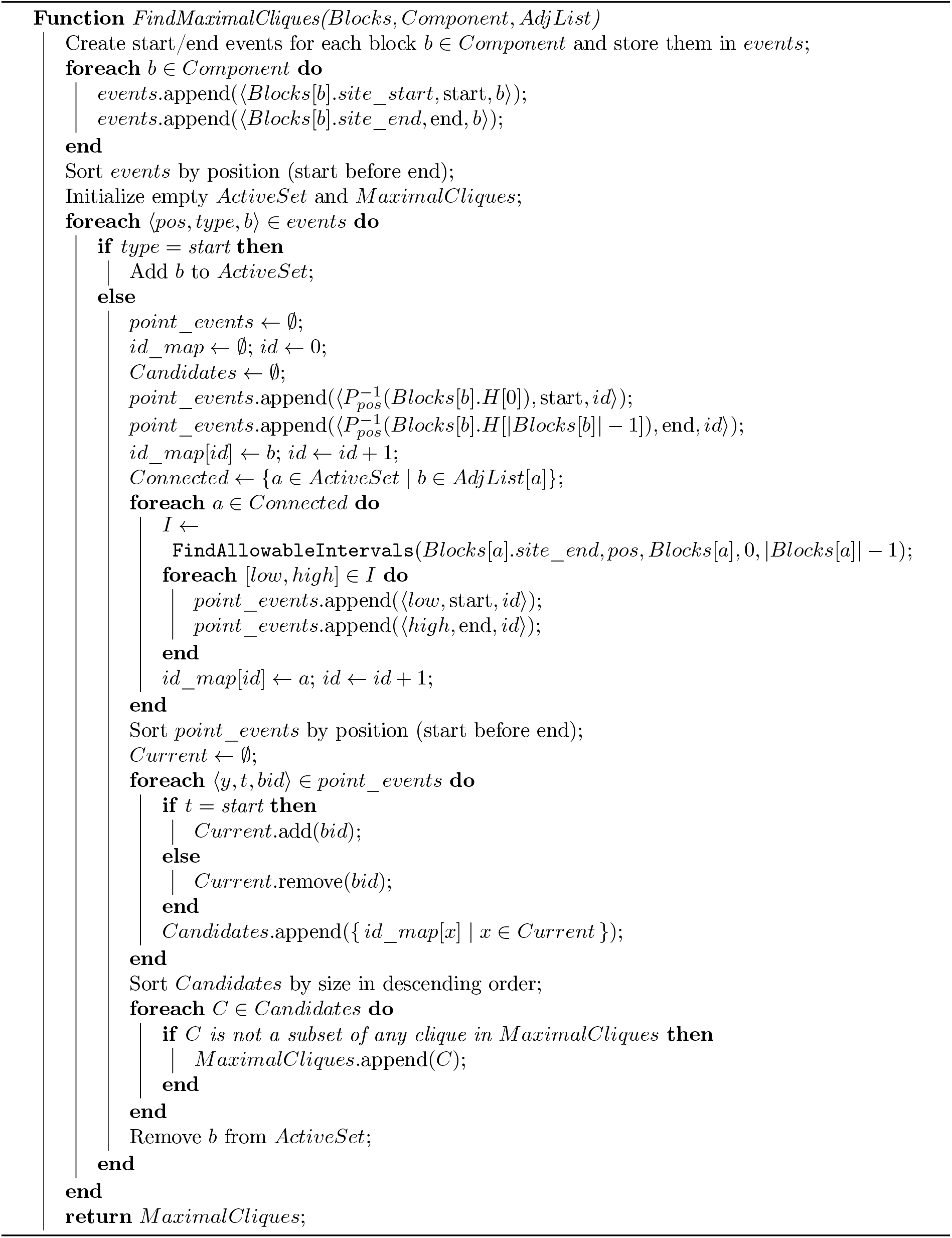

